# Prohibitin is a prognostic marker of relapse and therapeutic target to block chemotherapy resistance in Wilms tumor

**DOI:** 10.1101/508754

**Authors:** Michael V. Ortiz, Saima Ahmed, Melissa Burns, Anton G. Henssen, Travis J. Hollmann, Ian MacArthur, Shehana Gunasekera, Lyvia Gaewsky, Gary Bradwin, Jeremy Ryan, Anthony Letai, Ying He, Arlene Naranjo, Yueh-Yun Chi, Michael LaQuaglia, Todd Heaton, Paolo Cifani, Jeffrey S. Dome, Samantha Gadd, Elizabeth Perlman, Elizabeth Mullen, Hanno Steen, Alex Kentsis

## Abstract

Wilms tumor (WT) is the most common childhood kidney cancer. To improve risk stratification and identify novel therapeutic targets for patients with WT, we used high-resolution mass spectrometry proteomics to identify urine tumor markers associated with WT relapse. We determined urine proteomes at diagnosis of 49 patients with WT, non-WT renal tumors, and age-matched controls, leading to the quantitation of 6,520 urine proteins. Supervised analysis revealed specific urine markers of renal rhabdoid tumors, kidney clear cell sarcomas, renal cell carcinomas, as well as those detected in cured and relapsed WT. In particular, urine prohibitin was significantly elevated at diagnosis in patients with relapsed as compared to cured WT. In a validation cohort of 139 patients, a specific urine prohibitin enzyme-linked immunosorbent assay demonstrated that prohibitin concentrations greater than 998 ng/mL at diagnosis were significantly associated with ultimate WT relapse. Immunohistochemical analysis revealed that prohibitin was highly expressed in primary WT specimens and associated with disease stage. Using functional genetic experiments, we found that prohibitin was required for the growth and survival of WT cells. Overexpression of prohibitin was sufficient to block intrinsic mitochondrial apoptosis and to cause resistance to diverse chemotherapy drugs, at least in part by dysregulating factors that control apoptotic cytochrome c release from mitochondrial cristae. Thus, urine prohibitin may improve therapy stratification, non-invasive monitoring of treatment response and early disease detection. In addition, therapeutic targeting of chemotherapy resistance induced by prohibitin dysregulation may offer improved therapies for patients with Wilms and other relapsed or refractory tumors.

## Introduction

Wilms tumor (WT) is the most common kidney tumor in children. With current stratification and a combination of surgery, chemotherapy and radiotherapy, more than 90% of patients with low risk disease can now be cured. ^1^ However, treatment of patients with advanced, anaplastic, or relapsed disease remains challenging, with inadequate curative therapies and substantial long-term effects. ^1,2^

While anaplastic histology WT, most frequently due to inactivating mutations of *TP53*, is associated with unacceptably poor survival, distinct subsets of patients with favorable histology WT also suffer disease relapse. ^1,3,4^ Loss of heterozygosity (LOH) at 1p and 16q was found to be associated with inferior prognosis of favorable histology WT. ^4,5^ This led to new clinical trials to investigate whether intensification of therapy for patients with tumor LOH of 1p and 16q could be used to improve survival. However, additional biomarkers of adverse prognosis, therapy resistance and disease relapse will be needed for improved stratification of existing therapies and development of new therapies that are precise, curative, and safe. For renal tumors in particular, urine biomarkers offer the ability to monitor disease and therapy response non-invasively.

Neuron-specific enolase, basic fibroblast growth factor (bFGF), and hyaluronidase have been reported to be enriched in the urine of patients with WT. ^6–9^ In particular, elevation of urinary bFGF was correlated with WT disease stage. ^7^ Though its specificity and sensitivity were not sufficient to permit clinical use, continued elevation of urinary bFGF in a subset of WT patients who developed persistent or relapsed disease suggests that urine profiling may reveal prognostic biomarkers and improved therapeutic targets for WT and other kidney tumors.

We and others used high-resolution mass spectrometry proteomics to profile urine in order to identify improved disease biomarkers. ^10–13^ Here, we profiled urine proteomes of patients with diverse childhood kidney tumors as compared to age-matched controls. By comparing initial urine proteomes of patients who relapsed to those who were cured, we identified elevated urinary prohibitin (PHB) at diagnosis as a prognostic biomarker of relapse in favorable histology WT. Urinary PHB elevation was significantly associated with WT relapse in an independent patient cohort, with tumor PHB overexpression associated with WT disease stage. Using a battery of functional studies, we found that PHB overexpression regulates mitochondrial apoptosis and induces resistance to diverse chemotherapy drugs. These findings should enable improved WT therapy stratification and future strategies to overcome chemotherapy resistance to increase the cure rates of patients.

## Results

### Comparative urine proteomics of Wilms tumor, kidney rhabdoid tumor, kidney clear cell sarcoma, and renal cell carcinoma reveals new biomarkers

In previous studies, we optimized methods for the analysis of clinical urine proteomes, including protein isolation, fractionation, and high-resolution mass spectrometry. ^10–13^ For this study of childhood kidney tumors, we assembled a cohort of specimens collected at diagnosis from 49 patients, including 16 with favorable histology WT, 6 with rhabdoid tumor of the kidney (RTK), 9 with clear cell sarcoma of the kidney (CCSK), and 2 with renal cell carcinoma (RCC). For comparison, we included 16 age-matched control specimens collected from 10 healthy children and 6 children with acute abdominal pain who were evaluated as part of our prior study of acute appendicitis, and whose symptoms resolved spontaneously. ^11^ Mass spectrometric proteomic analysis of all specimens led to the identification of 6,520 urine proteins, detected with at least 2 unique peptides at the false discovery rate threshold of 1% (Figure 1A). As expected, supervised analysis of all kidney tumor specimens versus the age-matched controls revealed that urine tumor proteomes are dominated by markers of tissue injury and hematuria, consistent with kidney and blood vessel invasion (Supplemental Figure 1A).

**Figure 1.**
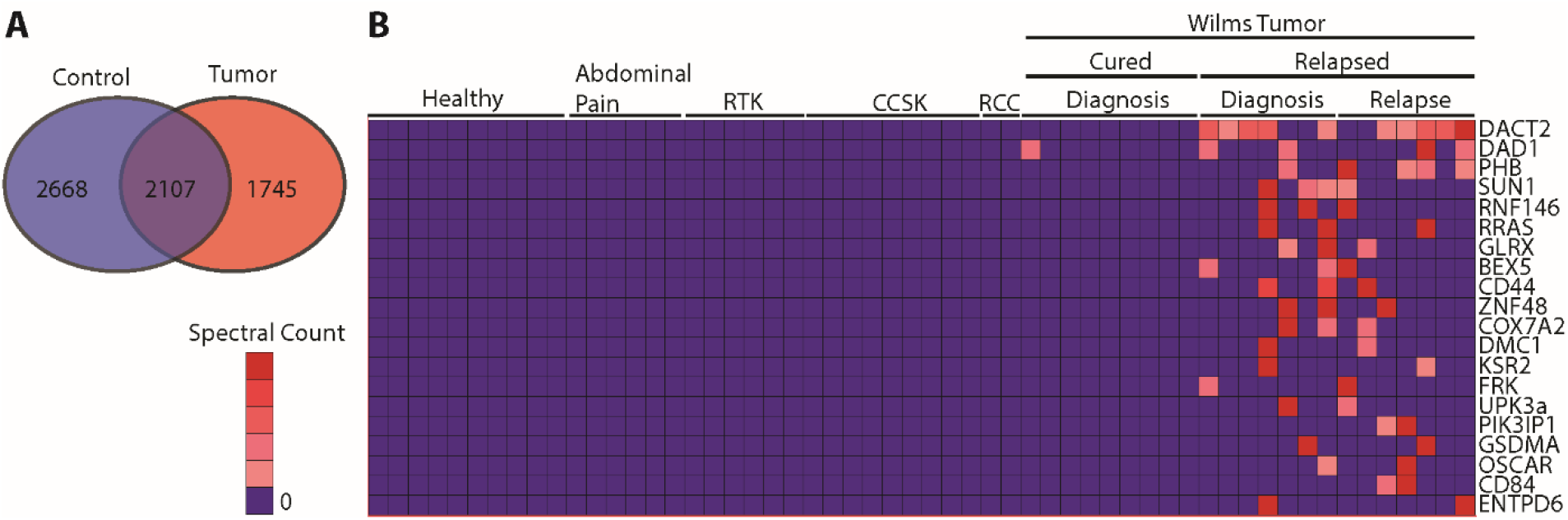
Using high accuracy mass spectrometry to profile the urine proteomes of childhood kidney tumors reveals markers of relapse and chemoresistance. **(A)** This venn diagram demonstrates the distribution of unique proteins identified in the children without renal tumors (Blue) compared to those with tumors (Red) (n = 6520 proteins). **(B)** The 20 proteins which were most highly enriched in Wilms tumors that relapsed are shown in this heat map (n=56 samples); RTK = Rhabdoid tumor of the kidney, CCSK = Clear cell sarcoma of the kidney, RCC = Renal cell carcinoma.

Importantly, supervised analysis of distinct tumor specimens revealed new tumor markers, such as PGBD5 (Supplemental Figure 1B), which we recently validated as an oncogenic DNA transposase and therapeutic target in rhabdoid tumors. ^14–17^ In the case of favorable histology WT, we identified the most abundant proteins specifically detected in WT as compared to other non-WT kidney tumor urine specimens (Supplemental Figures 1B-E). Finally, we stratified WT patients based on clinical outcome and identified urine proteins detected specifically in cases of relapsed favorable histology WT as compared to favorable histology WT (Figure 1B). These included β-catenin antagonist DACT2, mitochondrial regulators DAD1 and PHB, among others. Thus, comparative urine proteomics can be used to identify urine biomarkers, including new tumor markers that may represent improved biomarkers and therapeutic targets.

### Urine prohibitin is a prognostic marker of Wilms tumor relapse

To validate their prognostic significance and identify improved therapeutic targets of relapsed WT, we assembled an independent cohort of 139 specimens, including 99 favorable histology WT specimens, and 40 age-matched controls (Supplemental Table 1). First, we used enzyme-linked immunosorbent assay (ELISA) to measure protein concentration of candidate WT markers in clinical urine specimens. We used commercially available antibodies measure the top 3 urine markers enriched in patients with relapsed WT. While we could not develop specific ELISAs for DACT2 and DAD1 due to the limitations of commercially available antibodies, we confirmed that ELISA for PHB provided accurate measurements of urine PHB, with a linear signal response in the ng/mL range, as determined using purified recombinant PHB (Figure 2A). Thus, we measured the urine concentration of PHB, as compared to urine creatinine as a control for overall urine concentration (Supplemental Figure 2). We found that urine PHB was significantly enriched in diagnostic urine specimens from patients who ultimately relapsed (median 1672 ng/mL), as compared to those from patients who were ultimately cured (median 131 ng/mL) or age-matched controls (median 218 ng/mL). Using logistic regression, we determined that the PHB urine concentration of 998 ng/mL was significantly associated with WT relapse (odds ratio = 153.3, Figure 2B). Both urine PHB concentration and urine PHB concentration normalized to urine creatinine (Cr) were statistically significant predictors of relapse, independent of stage and therapy (Figure 2, Supplemental Figure 2). Similarly, receiver operating characteristic (ROC) curve analysis showed that urine PHB in patients with WT relapse had the prognostic area under the curve (AUC) of 0.78 (95% confidence interval 0.68-1.0; Figure 2C). Almost all patients with elevated urine PHB at diagnosis developed WT relapse within 2 years after therapy (Figure 2D). We confirmed that urine PHB elevation was not due to urine concentration, as evident by the lack of statistically significant differences in urine creatinine concentration (Supplemental Figure 2A-C). We found that urine PHB was significantly elevated in patients with abdominal as compared to lung WT relapse (Figure 2E). Indeed, urine PHB exhibited a near perfect prognostic performance for abdominal WT relapse, with the ROC AUC of 0.96 (95% confidence interval 0.91-1.0; Figure 2F). In all, these results indicate that urine PHB is a significant prognostic marker of WT relapse.

**Figure 2.**
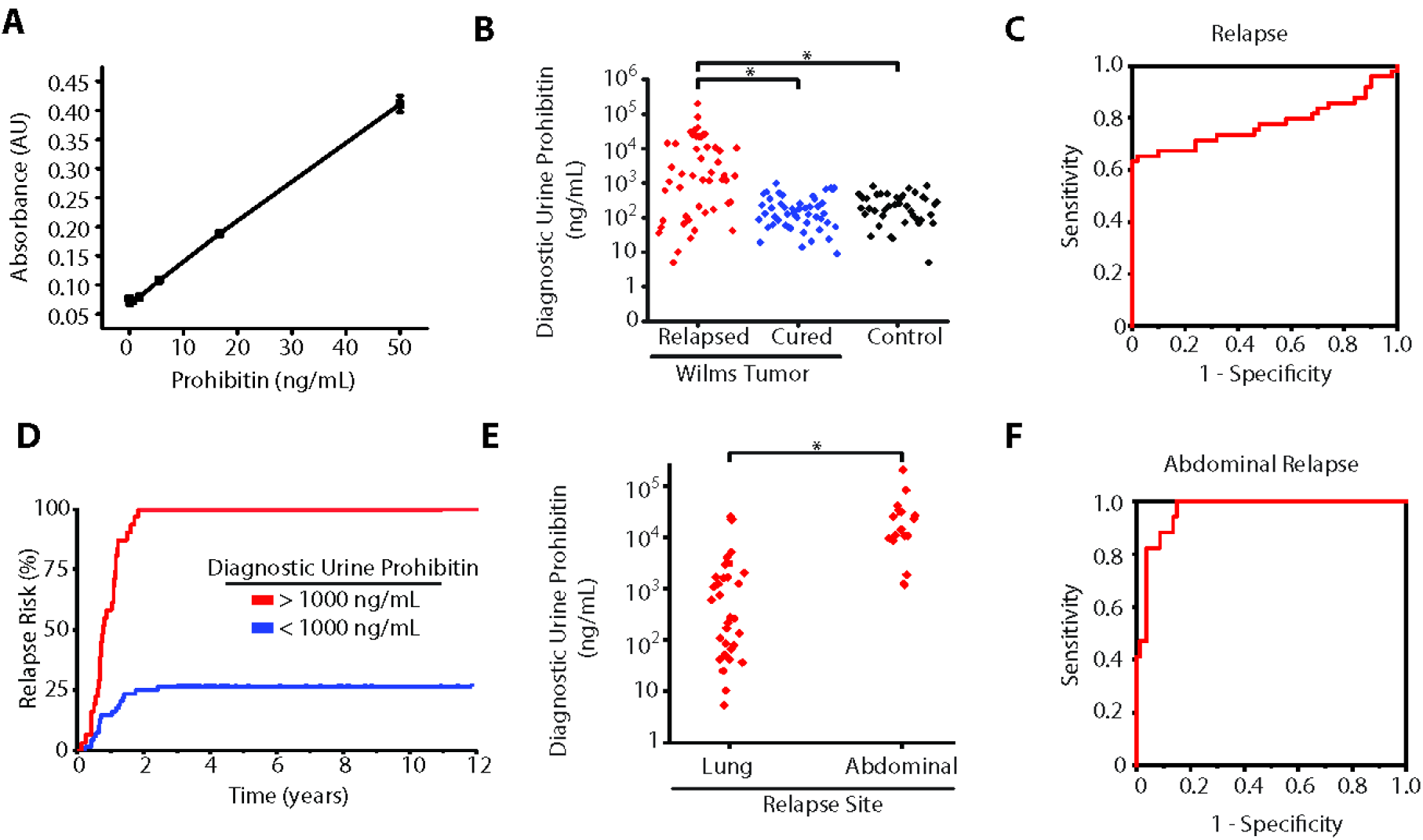
Elevated urine prohibitin at diagnosis is a specific biomarker of relapse in favorable histology Wilms tumors. (**A**) Enzyme linked immunosorbance assay (ELISA) comparing known prohibitin levels (ng/mL) with measured absorbance via ELISA. (**B**) Diagnostic urine prohibitin levels (ng/mL) in favorable histology Wilms tumor patients who relapsed (Red, n = 49) are compared with those who were cured (Blue, n = 50) and normal controls (Black, n = 40). Odds ratio of relapse for patients with diagnostic urine prohibitin > 1000 ng/mL = 153 (95% CI = 19.6 – 1000). (**C**) A receiver operating characteristic curve demonstrates the prognostic power of diagnostic urine prohibitin to predict relapse in favorable histology Wilms tumors at different sensitivity and specificity with an area under the curve of 0.77 (95% confidence interval 0.64-0.99). (**D**) Risk of relapse in patients with favorable histology Wilms tumors are stratified by those with a diagnostic urine prohibitin > 1000 ng/mL (Red, n = 31) compared with those with a diagnostic urine prohibitin < 1000 ng/mL (Blue, n = 68). (**E**) Diagnostic urine prohibitin levels in relapsed favorable histology Wilms tumor patients are stratified by site of relapse. (**F**) A receiver operating characteristic curve demonstrates the prognostic power of diagnostic urine prohibitin to predict abdominal relapse in favorable histology Wilms tumors at different sensitivity and specificity with an area under the curve of 0.99 (95% confidence interval 0.99-1.0).

### Prohibitin is overexpressed in Wilms tumor cells and correlates with tumor stage

To determine whether elevated PHB in WT patient urine samples was due to increased expression of PHB in WT cells, we used immunohistochemistry (IHC) using formalin-fixed paraffin embedded primary patient WT specimens, as compared to adjacent normal kidney tissue. We found that PHB was highly expressed in both favorable histology WT (Figure 3B and 3E), as well as diffusely anaplastic WT (Figure 3C and 3F). In agreement with prior studies, we also observed PHB expression in normal kidney tubules (Figure 3A and 3D). ^18^ We assembled a cohort of 59 primary favorable histology WT specimens and 10 control non-WT benign and malignant renal samples, uniformly stained for PHB (Supplemental Table 2). We scored IHC PHB expression in a blinded manner on the 0 to 3+ scale. Representative images of score levels are shown in Supplemental Figure 3. Notably, all Wilms tumors expressed PHB from 1+ to 3+ (Supplemental Figures 3D-F). Consistent with the specific detection of PHB in WT but not other kidney tumors (Figure 1B), we found no detectable PHB expression in pre-malignant nephrogenic rests (Supplemental Figure 3A), embryonal rhabdomyosarcoma (Supplemental Figure 3B), and clear cell sarcoma of the kidney (Supplemental Figure 3C). We found that PHB expression in WT correlated with higher tumor stage and increased percentage of tumors with higher PHB expression (Figure 3G).

**Figure 3.**
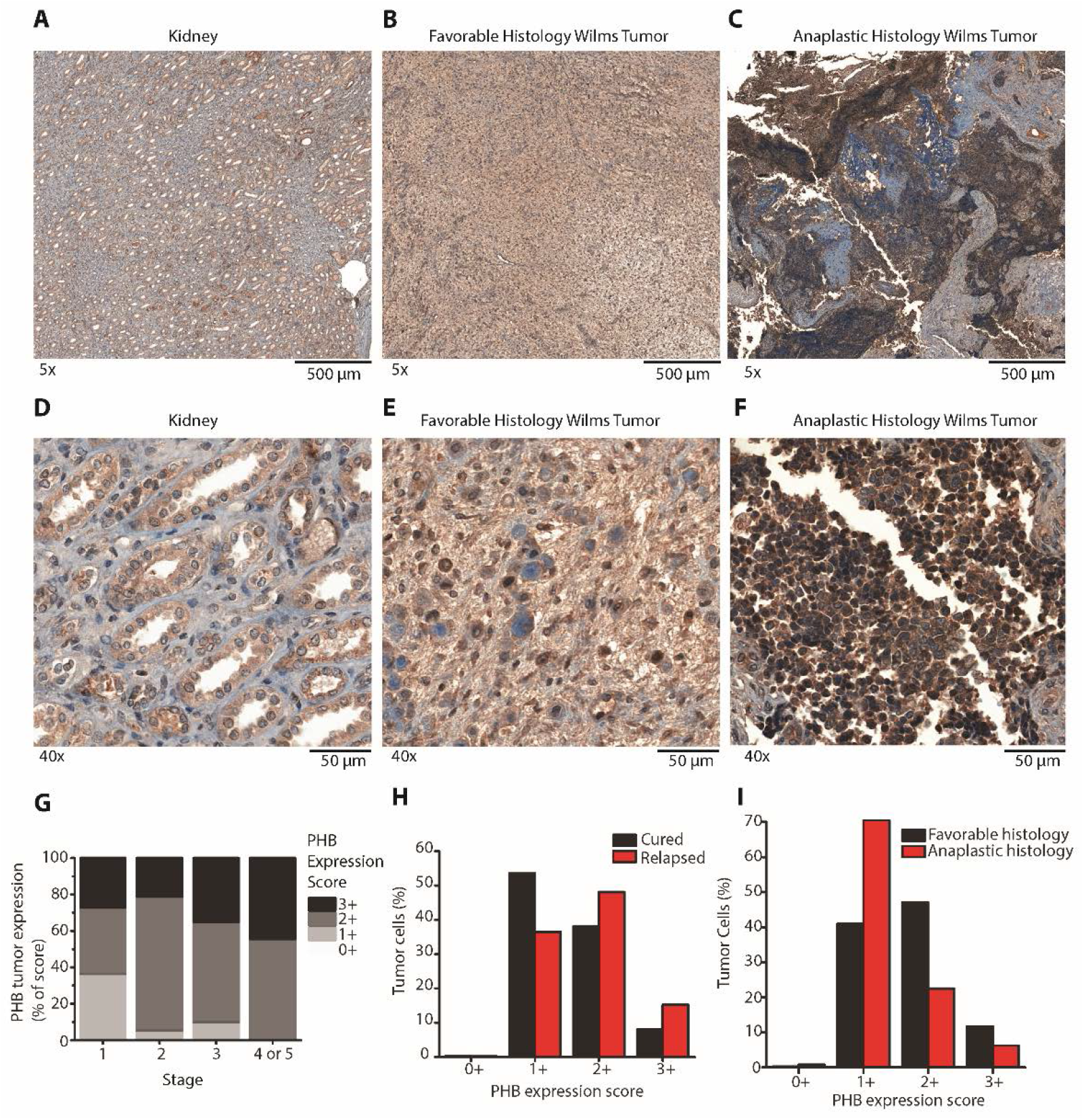
Prohibitin is highly expressed in primary Wilms tumor samples. (**A-F**). PHB immunohistochemical (IHC) staining was performed on formalin fixed paraffin embedded primary Wilms tumor samples and compared with adjacent normal kidneys. A-C are 5x magnified and E-F are 40x magnified. A and D include normal kidney. B and E include favorable histology Wilms tumors. C and F include anaplastic histology Wilms tumors. (**G**). IHC was performed on a tissue microarray containing 59 primary WT samples and graded from 0+ to 3+ in a blinded manner. Quantification of IHC from the tissue microarray is shown and stratified by initial tumor stage. (**H-I**). A second cohort of 38 primary WT patients was assessed (15 cured, 16 relapsed, 7 no information; 24 favorable, 14 anaplastic WT). The expression of PHB was evaluated on a cell-by-cell basis using Halo imaging analysis software. 24,862,509 cells in total were counted and scored from 0+ to 3+ based on PHB expression. In panel H, PHB expression in the cells of Wilms tumors that ultimately relapsed are compared with those that were cured. Whereas in panel I, PHB expression in the cells of Wilms tumors that had favorable histology are compared with those with anaplastic histology.

We further validated PHB expression using quantitative image densitometry in a second cohort of 38 primary WT patients, including both favorable histology and anaplastic WT (Supplemental Table 4). We observed increased PHB expression in favorable histology WT cells of specimens that ultimately relapsed, as compared with those that were cured with surgery and chemotherapy (Figure 3H). On a per-cell-basis, PHB expression was increased in favorable histology WT as compared to anaplastic histology WT (Figure 3I). We did not observe a statistically significant correlation between *PHB* mRNA expression and *TP53* mutations in diffusely anaplastic WT samples (Supplemental Figure 4). Likewise, we did not find recurrent mutations or amplification of *PHB* in a recently analyzed cohort of 117 patient WT specimens. ^19^ Though we found that cellular PHB expression was similar between WT and some normal kidney cells (Supplemental Figure 5A), we found that WT cells were 12% smaller, with 27% smaller cytoplasm and 11% larger nuclei, as compared to normal kidney cells (Supplemental Figure 5B). These findings suggest that WT cells overexpress PHB in specific subcellular compartments, presumably through post-transcriptional mechanisms.

**Figure 4.**
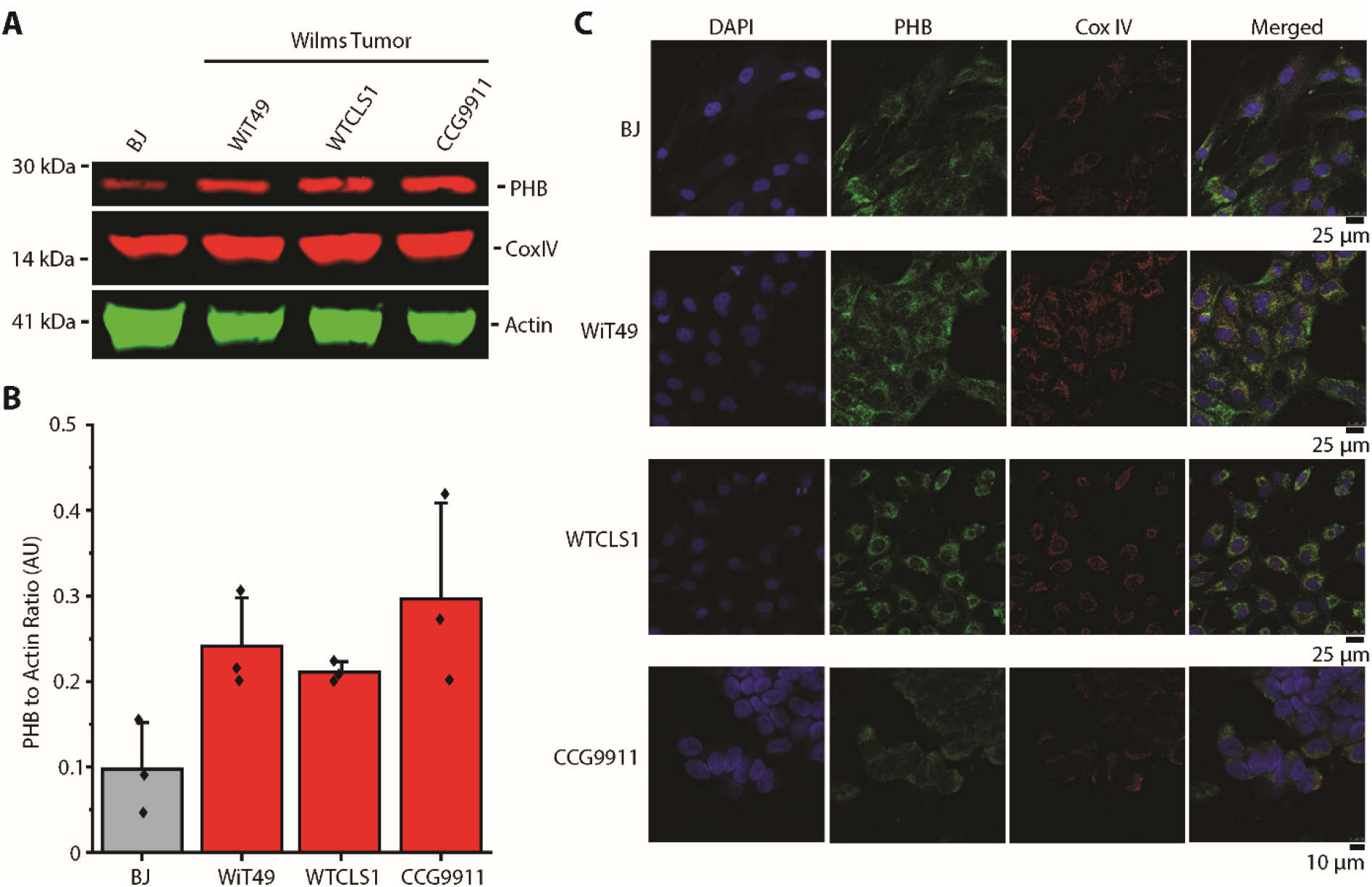
Prohibitin is an abundant mitochondrial protein in Wilms tumor cell lines. (**A**). Our *in vitro* studies of PHB included a control fibroblast cell line (BJ) as well as three Wilms tumor cell lines (WiT49, WTCLS1, and CCG9911). Endogenous PHB expression in the Wilms tumor cell line is shown with Actin as a loading control and CoxIV as a mitochondrial loading control. (**B**). PHB expression is compared in the different cell lines normalized to Actin in western blot triplicates. (**C**). Confocal imaging demonstrates colocalization of PHB (Green) with the inner mitochondrial membrane marker CoxIV (Red) but not the nuclear marker Dapi (Blue) in paraformaldehyde fixed cells.

### PHB is required for the growth and survival of Wilms tumor cells by regulating mitochondrial functions

Relative overexpression of PHB in WT cells suggests that PHB may contribute to their growth and survival. To assess this, we examined PHB expression and subcellular localization in WT cell lines WiT49, WTCLS1, and CCG9911, as compared to immortalized human BJ fibroblasts. Using quantitative fluorescent Western blot immunoassays, we found that PHB is more highly expressed in WT cell lines, as compared to BJ fibroblasts (Figure 4A). Normalized to cellular actin, PHB expression was an average 2.9, 2.7, and 3.3-fold higher in the WiT49, WTCLS1, and CCG9911 cells, respectively, as compared to normal BJ fibroblasts (Figure 4B).

Relative overexpression of PHB in WT cells and their relatively smaller size as compared to normal cells suggest that tumor PHB overexpression may be due to its increased expression in specific subcellular compartments, such as mitochondria. ^20^ Compelled by the finding of PHB staining in the cytoplasm of WT cells (Figure 3), we used confocal immunofluorescence microscopy to define the subcellular localization of PHB in WT cell lines. We observed that most of cellular PHB co-localized with the specific mitochondrial inner membrane marker CoxIV (Figure 4C). These findings are consistent with prior studies that identified PHB heterodimerization with PHB2 in association with the inner mitochondrial membrane to regulate mitochondrial morphogenesis. ^21–26^ Thus, we reasoned that PHB overexpression may contribute to the growth or survival of WT cells. To test this hypothesis, we used three independent short hairpin RNA (shRNA) interference lentiviral constructs to deplete PHB in WT cell lines, as compared to the control shRNA targeting the green fluorescent protein (GFP) which is not expressed (Figure 5 A, B, and C). We confirmed PHB depletion using Western immunoblotting (Figure 5 A, B, and C). Consistently, cells depleted of PHB exhibited significantly decreased proliferation, as compared to wild-type cells or those expressing the control GFP-targeting shRNA (Figure 5 D, E, and F).

**Figure 5.**
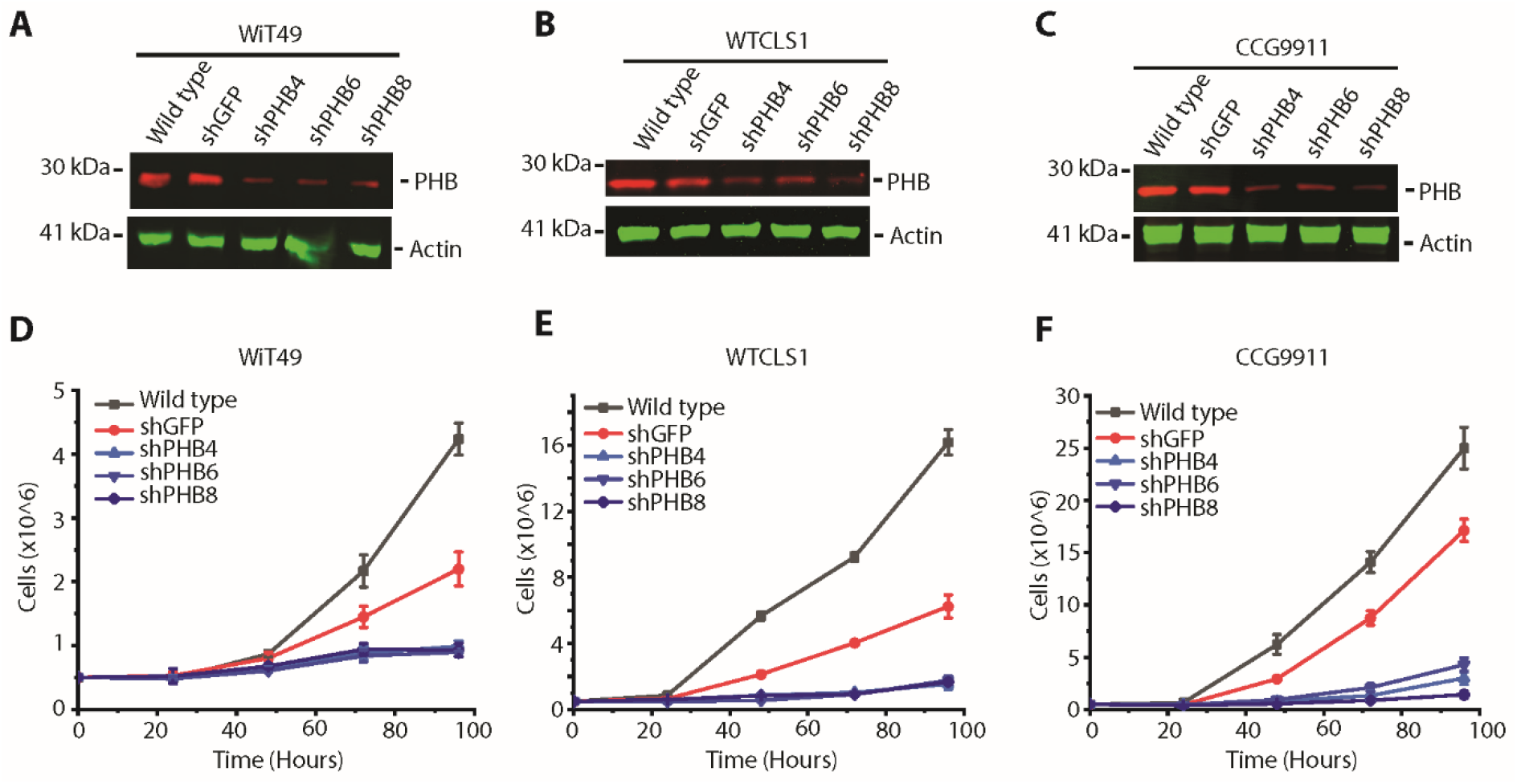
Prohibitin is required for Wilms tumor growth and survival. (**A-C**). Western blots of wild type Wilms tumor cells (A, WiT49; B, WTCLS1; C, CCG9911) as well as a nontargeting shGFP and three different shRNA hairpins targeting the PHB 3’UTR (shPHB4) and CDS (shPHB6 and shPHB8). Cox IV is used for mitochondrial loading control and Actin is used for whole cell loading control. (**D-F**). Cell growth over time in Wilms tumor cells (D, WiT49; E, WTCLS1; F, CCG9911) which are wild type (Black), compared with those transduced with non-targeting shGFP (Red), and the three PHB targeting shRNA (Blue).

PHB has been reported to regulate mitochondrial morphology by interacting with the OPA1 GTPase, and the YMEL1 and OMA1 proteases that proteolytically process OPA1 that can control the release of cytochrome c from mitochondrial cristae during apoptosis. ^27 28,29^ We found that WT cells have elevated levels of OMA1 and variable levels of YME1L relative to BJ fibroblasts (Figure 6A). Consistent with the putative interaction of PHB with OPA1, depletion of PHB was associated with apparent reduction of OMA1 in all WT cell lines tested, but not in control cells transduced with GFP-targeting controls (Figure 6B-D). Since YMEL1 can proteolytically cleave OPA1 ^30,31^, we analyzed apparent OPA1 isoforms by Western immunoblotting. We found that cells with elevated endogenous YME1L, such as WTCLS1, depletion of PHB was associated with a reduction of YME1L and the S4 isoform of OPA1 (Figure 6D). Conversely, in cells with relatively low endogenous YME1L, such as WiT49 and CCG9911, depletion of PHB increased YME1L and the S4 OPA1 isoform (Figures 6B and C). In all, these findings indicate that PHB directly or indirectly interacts with the mitochondrial intermembrane proteases OMA1 and YME1L that cooperatively process OPA1.

**Figure 6.**
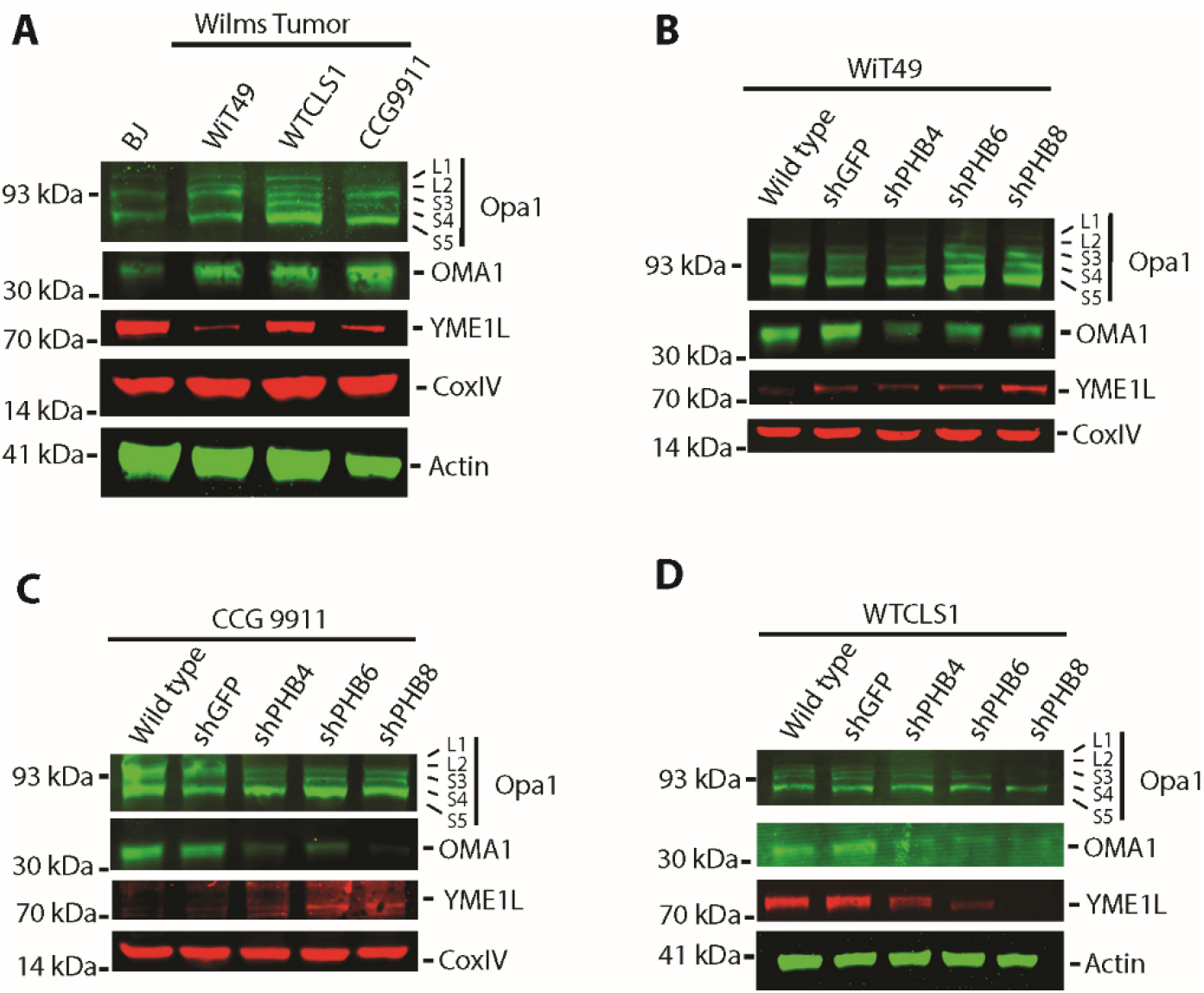
Depletion of prohibitin results in alterations in mitochondrial intermembrane proteases and structural proteins involved in apoptosis and mitochondrial morphogenesis. (**A**). Western blot of endogenous expression of Opa1, OMA1, and YME1L with Cox IV and Actin as mitochondrial and whole cellular loading controls, respectively. (**B-D**). Western blots of Opa1, OMA1, and YME1L in Wilms tumor cells (B, WiT49; C, WTCLS1; D, CCG9911) comparing wild type cells with nontargeting shGFP and three different shRNA hairpins targeting the PHB 3’UTR (shPHB4) and CDS (shPHB6 and shPHB8). Loading controls as shown.

### PHB overexpression causes resistance to mitochondrial apoptosis and diverse chemotherapy drugs

PHB-mediated control of OPA1 processing that can regulate apoptotic cytochrome c release raises the possibility that PHB overexpression in WT may impair mitochondrial apoptosis and cause chemotherapy resistance. To test this hypothesis, we ectopically overexpressed PHB in WT cell lines WTCLS1 and WiT49 as well as BJ fibroblasts using lentiviral transduction and confirmed transgene expression by Western immunoblotting of PHB and its V5 epitope tag in two independent clones (Figure 7). Even though we found relatively modest overexpression of PHB as compared to its endogenous levels, all PHB-overexpressing cells exhibited significantly increased resistance to vincristine, doxorubicin and dactinomycin, as compared to wild-type cells or control cells transduced with empty vectors (Figure 8). This effect was more pronounced in BJ fibroblasts and WTCLS1 WT cells, as compared to WiT49 WT cells, consistent with the relatively higher levels of endogenous PHB in WiT49 cells (Figure 4).

**Figure 7.**
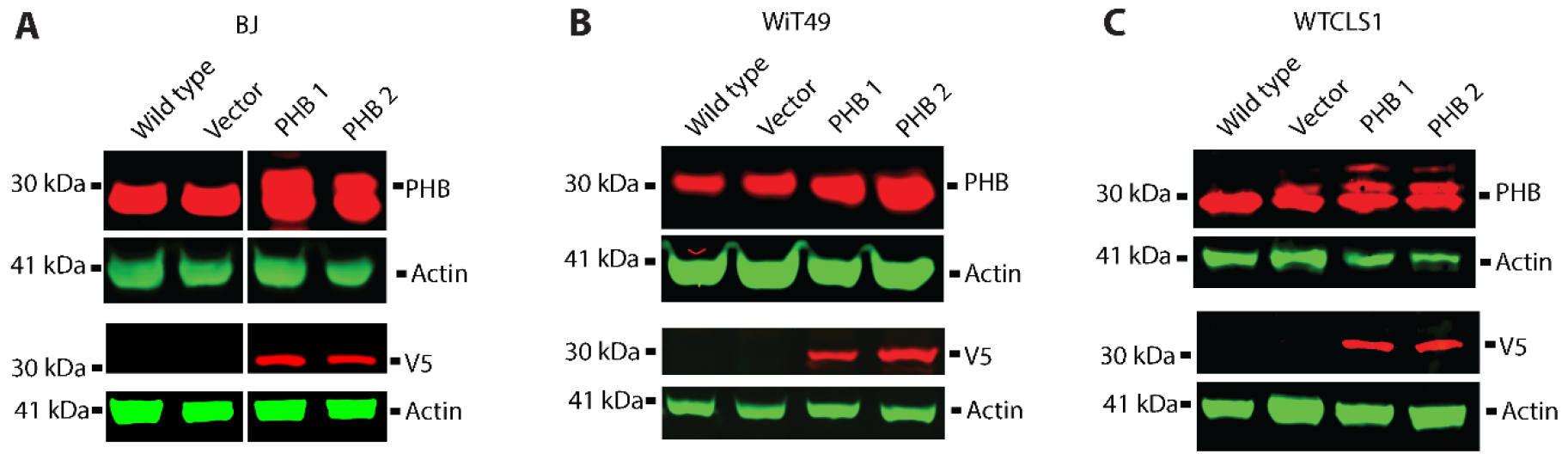
Overexpression of prohibitin in Wilms tumor and control cell lines. (**A-C**). Western blots of PHB and V5 in control (A, BJ) and Wilms tumor cells (B, WiT49; C, WTCSL1) comparing wild type cells and those transduced with an empty vector with two different clones transduced with a PHB expressing vector containing a V5 tag.

**Figure 8.**
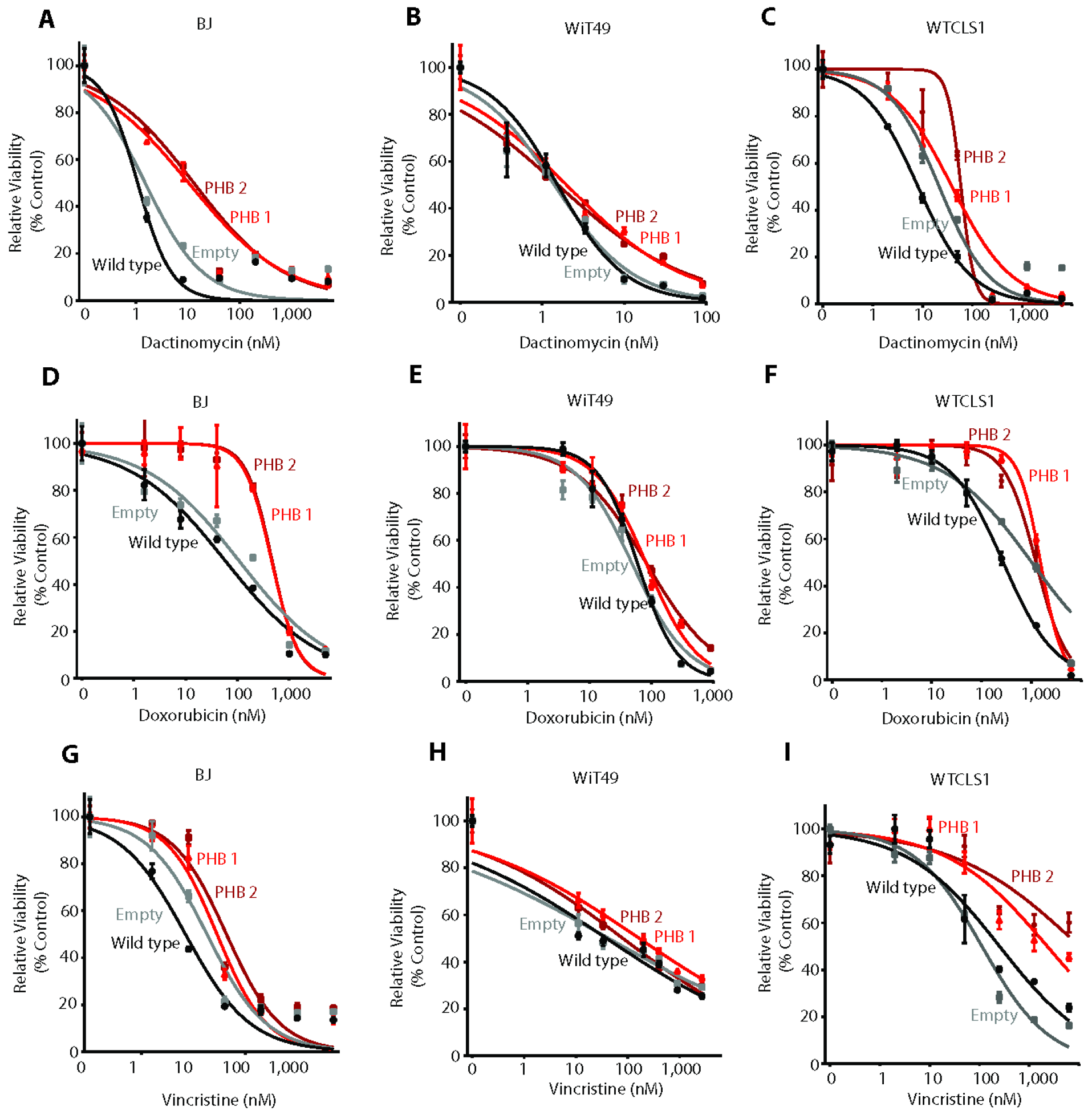
Overexpression of prohibitin results in resistance to chemotherapy in both Wilms tumor and control cells. (**A-I**). Dose response curve of BJ (D, G, J), WiT49 (E, H, K), and WTCLS1 (F, I, L) cells treated with Dactinomycin (D-F), Doxorubicin (G-I), or Vincristine (J-L) for 72 hrs comparing wild type cells (Black), empty vector transduced cells (Gray), as well as PHB transduced cells (Red, Dark Red).

Overexpression of PHB in patient Wilms tumors and urine, its requirement for enhanced WT cell growth, control of OPA1 and other factors that can regulate mitochondrial cytochrome c release, and sufficiency to cause resistance to chemotherapy drugs with diverse mechanisms of action suggest that PHB overexpression may contribute to WT therapy failure and relapse by blocking intrinsic mitochondrial apoptosis. To test this prediction, we used BH3 profiling, a dynamic assay of mitochondrial apoptotic function, of WiT49 WT and normal BJ cells (Figure 9, Supplemental Figure 6), as optimized to specifically measure mitochondrial cytochrome c release using flow cytometry. ^32^ Upon activation of mitochondrial apoptosis, we observed that PHB overexpression caused increased resistance to multiple different synthetic BH3 apoptotic activators, including PUMA, BAD, and BID (Figure 9, Supplemental Figure 6).

**Figure 9.**
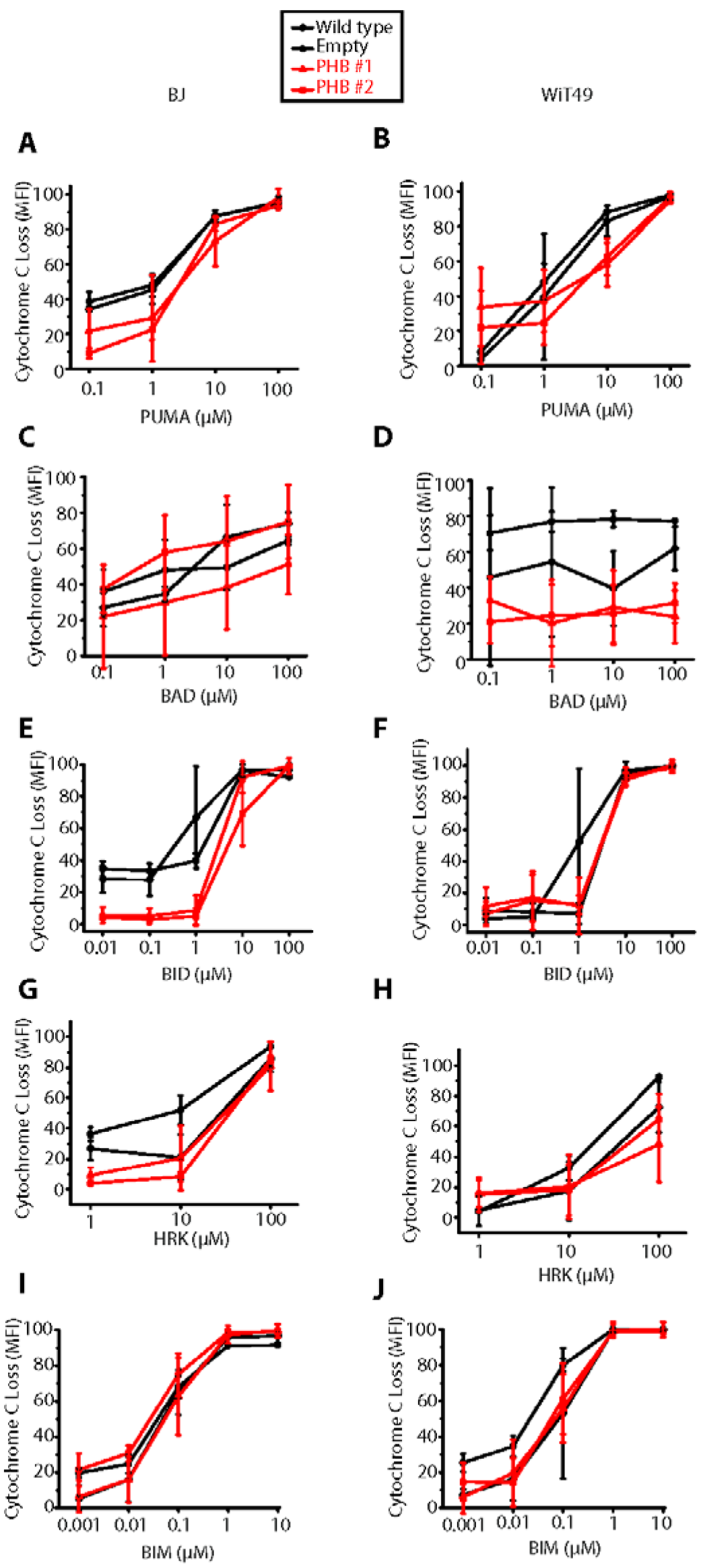
BH3 profiling reveals globally decreased apoptotic priming in response to PHB overexpression. (**A-J**). Cytochrome c loss in response to treatment with different pro-apoptotic peptides comparing Wild type (Black Diamond), with Empty (Black Circles), and two PHB overexpressing cell lines (Red Triangle, Red Square) in both BJ control fibroblasts and WiT49 Wilms tumors.

To further elucidate mitochondrial abnormalities in WT, we used transmission electron microscopy to determine the mitochondrial structure of newly diagnosed favorable histology WT immediately following nephrectomy. We observed that WT cells exhibited smaller mitochondria with blunted cristae and reduced matrix density, as compared to normal kidney tissue (Figure 10A-C). This is consistent with abnormal mitochondrial fission in WT, in agreement with aberrant mitochondrial morphogenesis induced by PHB overexpression, and prior ultrastructural studies of WT. ^33^ Thus, PHB overexpression causes intrinsic mitochondrial apoptosis resistance, thereby promoting chemotherapy resistance and WT therapy failure.

**Figure 10.**
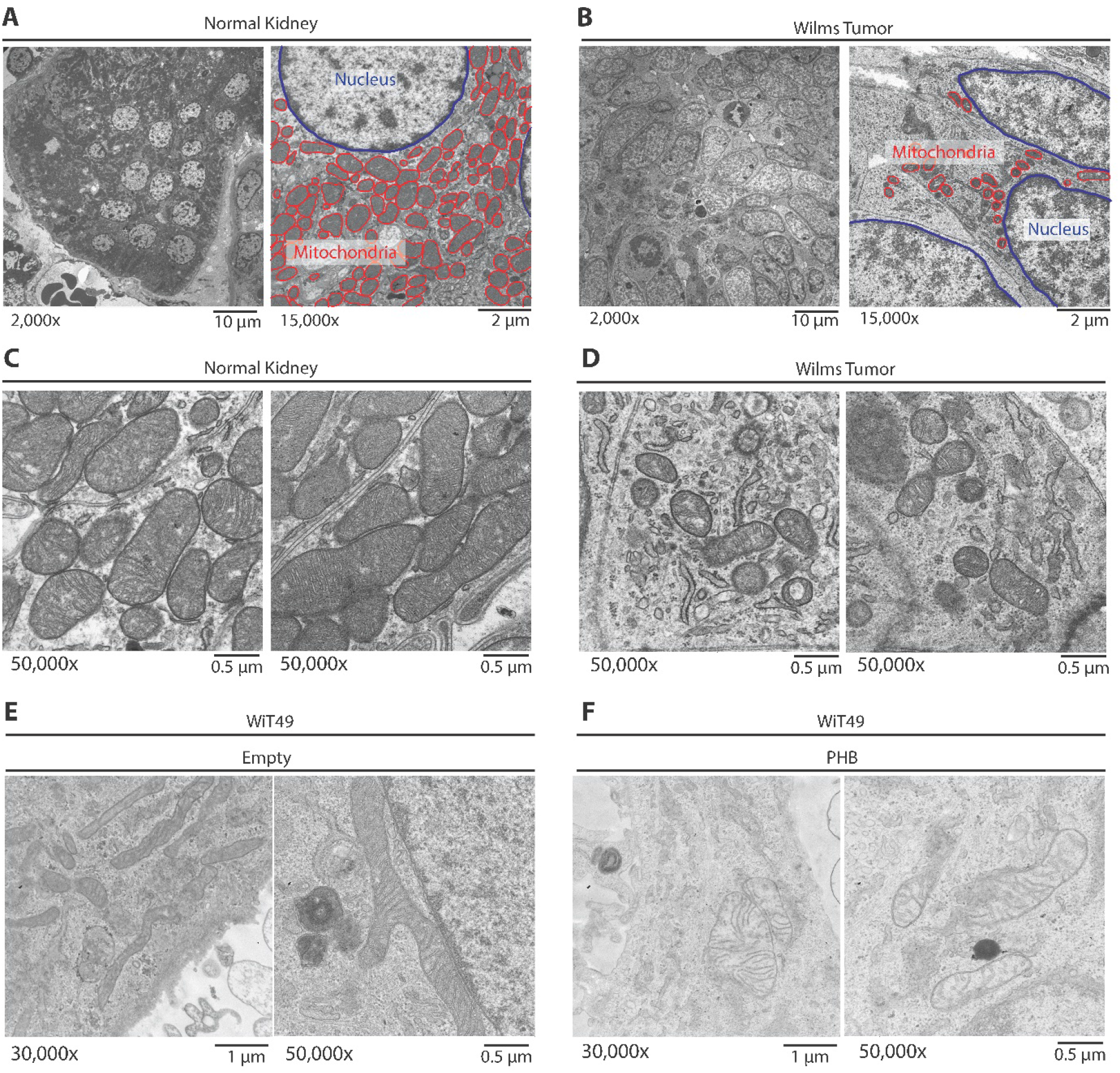
Primary Wilms tumor samples exhibit fewer mitochondria and predominantly a fission phenotype, as compared with adjacent normal kidneys. (**A-D**). Untreated favorable histology Wilms tumor was immediately fixed following nephrectomy and imaged using a transmission electron microscope to evaluate mitochondrial morphology. The normal kidney (Panel A, 2,000x magnification; Panel C, 50,000x magnification) is compared with the Wilms tumor (Panel B, 2,000x magnification; Panel D, 50,000x magnification). Panels A and B also highlight at 15,000x the size and number of the mitochondria (red) as compared with the nucleus (blue). (**E-F**). Characteristic mitochondria of a Wilms tumor cells (WiT49) treated with an empty vector (**E**) compared with a PHB overexpressing vector (F).

## Discussion

Pathogenesis of childhood solid tumors remains poorly understood. ^15^ In particular, distinct renal tumors remain difficult to treat, with limited knowledge of prognostic biomarkers and therapeutic targets. In the case of Wilms tumors, the most common childhood kidney tumor, subsets of patients remain incurable, with no effective means to monitor therapy response, stratify existing therapies, and identify improved therapeutic targets. In this context, our findings are significant for several reasons. First, reported urine proteome profiles of diverse kidney tumors provide a rich source of tumor biomarkers and therapeutic targets. Indeed, our recent study of PGBD5, identified in the proteomic profiles of renal rhabdoid tumors, revealed an unanticipated mechanism of solid tumor pathogenesis and improved therapeutic targets. ^15–17^

In the case of Wilms tumors, we found that overexpression of PHB at diagnosis is a prognostic marker of WT relapse. Levels of urinary PHB above 998 ng/mL were found to be significantly associated with relapse in independent cohorts of WT patients (Figures 1 and 2). PHB overexpression was found to be elevated not only in urine but also in tumor tissue samples in both favorable and anaplastic histology WT, and primarily localized to the mitochondria (Figures 3 and 4). We found that PHB was required for the growth and survival of WT cells, at least in part by regulating mitochondrial function and apoptosis (Figures 5 and 6). Importantly, ectopic overexpression of PHB was sufficient to cause resistance to diverse chemotherapy agents used for clinical treatment of WT. PHB overexpression blocked cytochrome c release from mitochondria, an essential step in the initiation of intrinsic mitochondrial apoptosis (Figures 7–9), at least in part by dysregulating mitochondrial cristae (Figure 10). Thus, we conclude that PHB overexpression impairs mitochondrial morphogenesis and apoptotic priming, and causes chemotherapy resistance, as schematized in Figure 11, which can be monitored non-invasively by urine PHB measurements.

**Figure 11.**
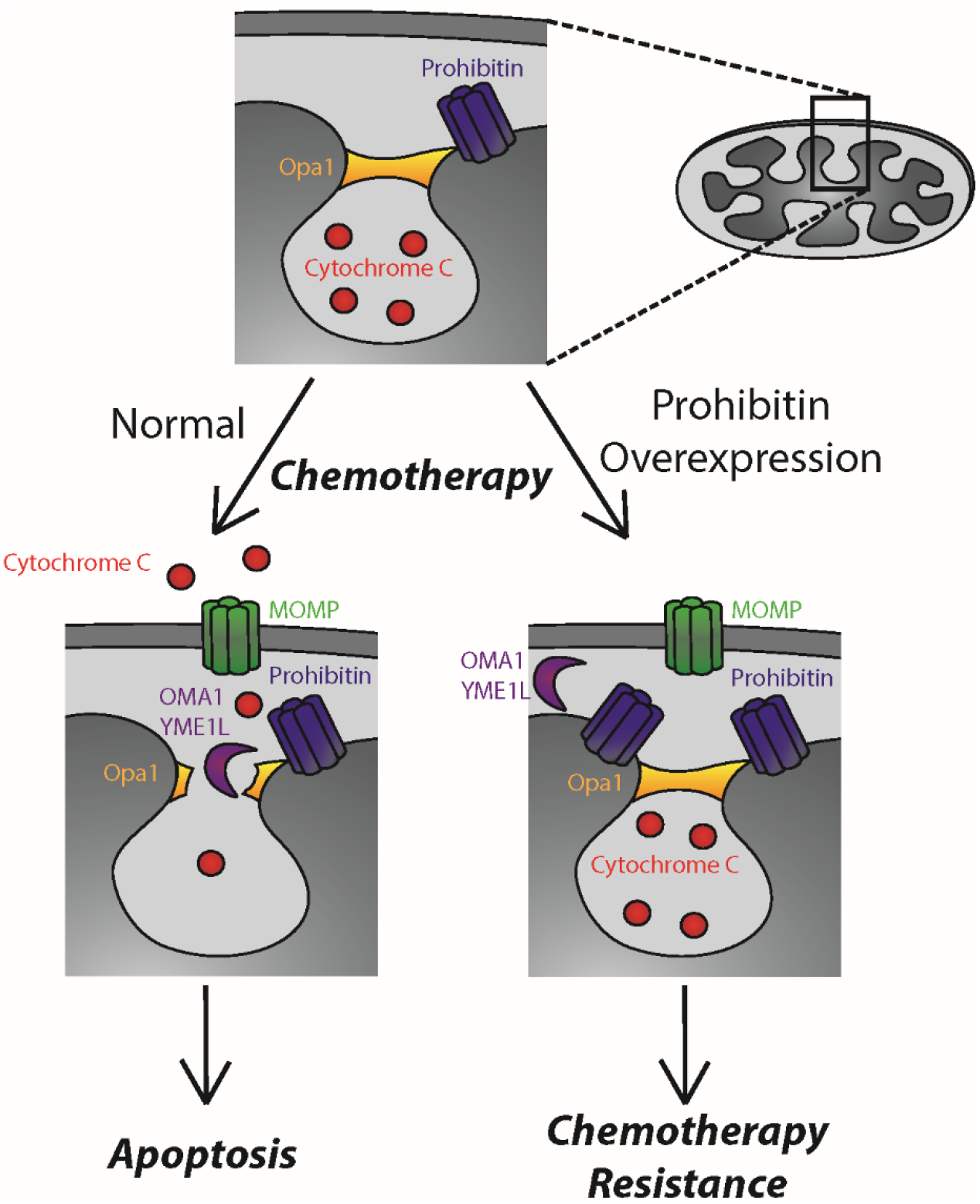
PHB overexpression leads to failure to release cytochrome c from cristae junctions despite the presence of a MOMP complex. Prohibitin (Blue) forms a complex along the mitochondrial inner membrane that interacts with several key regulators of apoptosis, including the Opa1 complex (Orange) located at cristae junctional openings. The majority of cytochrome c (Red) is sequestered within these cristae and in response to an apoptotic stimuli such as chemotherapy, the mitochondrial outer membrane pore (MOMP) complex consisting of BAX and BAK is inserted into the mitochondrial outer membrane. Upon proteolysis of the Opa1 complex, generally via OMA1 or other intermembrane proteases (Purple), Cytochrome c is released into the cytosol where it results in the apoptotic cascade. Our model proposes that due to the overexpression of PHB, in response to apoptotic stimuli, intermembrane proteases are no longer able to access the Opa1 complex thereby resulting in resistance to chemotherapy despite the presence of a MOMP pore in the outer mitochondrial membrane, ultimately resulting in failure to undergo apoptosis and chemotherapy resistance.

Overexpression of PHB has been described in other tumors, such as breast carcinomas, where somatic amplifications of the *PHB* gene have been reported in a subset of tumors. ^34–36^ We did not observe any cases of WT with *PHB* mutations or amplifications, suggesting that WT overexpression of PHB may involve post-transcriptional mechanisms. *PHB* expression has been found to be regulated by miR-27a in other tissues ^37–40^, and miR-27a is significantly downregulated in chemotherapy resistant blastemal Wilms tumors. ^41^ Since several miRNA processing genes are recurrently mutated in subsets of Wilms tumors, it is possible that PHB overexpression is caused by miRNA dysregulation, potentially including miR-27a. Future studies will be needed to determine the mechanisms of potential regulation of *PHB* expression by miRNAs and causes of its pathogenic overexpression. Likewise, additional studies will be needed to determine the prevalence and function of PHB in highly chemotherapy resistant anaplastic WT.

We found urine PHB to be elevated in the majority of favorable histology WT patients whose disease ultimately relapsed (Figure 2B). These diagnostic urine PHB levels were generally orders of magnitude higher in patients with ultimately relapsed WT as compared to cured patients (Figure 2B). However, in primary tumor tissue samples, PHB was significantly but modestly overexpressed in relapsed WT as compared to cured WT and normal kidney tissue (Figure 3, Supplemental Figure 5). Thus, urine PHB elevation may be due to increased WT cell invasion and PHB release into renal tubules, as suggested by the increased invasiveness of colorectal and lung carcinomas with PHB overexpression. ^42–44^ This possibility is further supported by the relatively higher frequency of abdominal WT relapse in patients with elevated urine PHB (Figure 2E). Additional studies will be needed to establish the mechanisms and significance of this possible phenomenon, and its relationship with abnormal mitochondrial structure and function induced by PHB overexpression. However, if PHB overexpression indeed causes increased WT cell invasion, then augmenting local control measures such as additional lymph node dissection and radiation therapy may be helpful.

Our findings implicate the interaction between PHB and the mitochondrial proteases OMA1 and YMEL1 in the dysregulation of mitochondrial cristae function and apoptotic cytochrome c release. It is possible that the ring complex formed by PHB and PHB2 affects the OPA1 GTPase that remodels mitochondrial cristae by sequestering the inner mitochondrial membrane OMA1 and YMEL1 proteases within microdomains which are unable to access OPA1. ^21,45^ As a result, PHB overexpression would impair the release of cytochrome c from mitochondrial cristae, blocking the initiation of mitochondrial apoptosis, as well as altering the dynamics of mitochondrial morphogenesis, which could further contribute to alterations in apoptosis. ^28–31,46^ In addition, although the PHB complex is associated with the inner mitochondrial membrane, it has been shown to interact with BAX and BAK that comprise the apoptotic MOMP pore. ^20,47^ Thus, in addition to modulating cytochrome c release at crista junctions, PHB overexpression may also impair the formation or function of the MOMP pore. Third, PHB overexpression may impair apoptosis because the properties of the mitochondrial outer membrane govern the ability of the predominantly cytosolic BAX to insert into the mitochondrial outer membrane as well as its ability for subsequent activation by BH3-only proteins. ^48,49^ Notably, this impaired BAX membrane insertion would be expected in smaller, hyper-fragmented mitochondria produced as a result of mitochondrial fission. ^48^ Lastly, PHB overexpression can also affect the energetic and metabolic mitochondrial functions, as well as its other cellular functions, such as regulation of cytosolic signaling. For example, PHB has been reported to interact with the mitochondrial respiratory machinery and thus it is also possible that PHB mediated overexpression co-opts mitochondrial respiration. ^26^ An analogous mechanism was described wherein inhibitory factor 1 mediated a decrease in OMA1 proteolysis and resulted in impaired Opa1 cleavage by maintaining ATP levels thereby reducing glutathione consumption and inactivating peroxiredoxin 3 during apoptosis. ^50^ Importantly, given that rocaglamides and aurilide can interact with PHB, it is possible that their derivatives can be developed to specifically interfere with the oncogenic functions of PHB in WT and other tumors. ^51–54^

In all, PHB-mediated evasion of apoptosis through dysregulation of mitochondrial cristae structure represents a previously unanticipated mechanism by which WT cells resist cell death, thereby causing therapy failure and WT relapse. Non-invasive monitoring of urine prohibitin offers an immediately accessible biomarker to identify and treat patients at risk for WT relapse, which should be investigated in future prospective clinical trials.

## Methods

### Specimens

Urine and tumor specimens were obtained from the Children’s Oncology Group and Memorial Sloan Kettering Cancer Center, as part of institutional review board-approved research studies, with informed subject consent.

### Mass Spectrometry

Urine specimens were fractionated and the protein composition of the fractions was identified by using high-resolution mass spectrometry as described previously ^13^. Briefly, individual five ml urine aliquots were fractionated using ultracentrifugation, protein precipitation, and denaturing polyacrylamide gel electrophoresis, and protein fractions were reduced, alkylated and trypsin digested. Urine protein fractions were subjected to liquid chromatography tandem mass spectrometry using a nanoflow HPLC system (Eksigent, Dublin, CA) coupled to the hybrid linear ion trap-Fourier transform ion cyclotron resonance mass spectrometer (LTQ FT Ultra, Thermo Scientific, Waltham, MA). Resultant spectra were processed to extract the 200 most intense peaks and searched against the human International Protein Index database (version 3.69) by using MASCOT version 2.1.04 (Matrix Science). Assessment of identification accuracy was carried out by searching a decoy database composed of reversed protein sequences of the target IPI database. Only proteins identified on the basis of two or more unique peptides were included in the analysis, at a false discovery rate of <1% at the peptide level. Mass spectrometry data are openly available at PeptideAtlas (http://www.peptideatlas.org/PASS/PASS00248).

### PHB ELISA

Enzyme linked immunosorbent assay (ELISA) against human PHB was constructed using the sandwich capture method. The capture antibody was a rabbit polyclonal antibody against the C-terminal domain and the detection antibody was a mouse monoclonal antibody paired with the capture antibody and labeled with biotin. The blocking buffer was composed of 1% bovine serum albumin in phosphate-buffered saline. Recombinant human prohibitin was produced in *E.coli* as a single, non-glycosylated His-tagged protein. Streptavidin-horseradish peroxidase with tetramethylbenzidine as a substrate was used for detection, as available from Novatein Biosciences (BG-HUM11702; Woburn, MA, United States).

### Immunohistochemistry and Immunofluorescence

Immunohistochemistry was performed using established methods as described previously. ^55^ Images were analyzed using the Halo imaging analysis software (Indica Labs; Corrales, NM, United States). 24,862,509 cells in total were counted and scored from 0+ to 3+ based on PHB expression. Confocal immunofluorescence microscopy was performed using cells plated in 4-well glass Millicell EZ slides fixed in 4% paraformaldehyde and washed with PBS prior to imaging. PHB and CoxIV antibodies used are listed in the Western Blotting section below, and were stained at 2.5 and 1.2 μg/mL, respectively. Images were acquired using the Leica TCS SP5 II (Leica Microsystems Inc., Buffalo Grove, IL, United States).

### Cell culture

HEK293T and BJ cells were obtained from the American Type Culture Collection (Manassas, VA, United States). WTCLS1 cells were obtained from the Leibniz Institute – Deutsche Sammlung von Mikroorganismen und Zellkulturen (Braunschweig, Germany). CCG-9911 cells were a gift from Benjamin Tycko at Columbia University (New York, New York, United States). WiT49 cells were a gift from Herman Yeger at University of Toronto (Toronto, Ontario, Canada). The identity of all cell lines was verified by STR analysis and lack of Mycoplasma contamination was confirmed by Genetica DNA Laboratories (Burlington, NC, United States). BJ cells were cultured in DMEM; WTCLS1 and CCG-9911 cells were cultured in IMDM; and WiT49 cells were cultured in DMEM F-12. All media included 10% fetal bovine serum, 100 U/mL penicillin, 100 μg/mL streptomycin, and 1% glutamine, as maintained in a humidified atmosphere at 37°C and 5% CO_2_.

### Plasmids

For RNA interference, pLKO.1 vectors targeting PHB were obtained from the RNAi Consortium: TRCN0000029204 (referred to as shPHB4), TRCN0000029206 (referred to as shPHB6), and TRCN0000029208 (referred to as shPHB8). PHB was overexpressed using pLX304 vector, as obtained from the DNASU repository. All plasmids were verified by restriction endonuclease mapping and Sanger sequencing, and are available from Addgene (https://www.addgene.org/AlexKentsis/).

### Lentivirus Transduction

Lentivirus particles were produced using HEK293T cells with the psPAX2 and pMD2.G packaging plasmids, as described previously. ^16^ Cell transduction was performed using the multiplicity of infection of 10 in the presence of 8 μg/ml hexadimethrine bromide. Transduced cells were selected using puromycin (3 μg/mL) for the pLKO.1 vectors or with blasticidin (3 μg/mL) for the pLX304 vectors. Transduced cells were cloned using limiting dilution.

### Western Blotting

Western blotting was carried out as described.^14^ PHB antibody used was a rabbit polyclonal antibody (1:1000, H-80) from Santa Cruz Biotechnology (Dallas, TX, United States). OMA1 antibody used was a mouse monoclonal antibody (1:1000, H-11) from Santa Cruz Biotechnology (Dallas, TX, United States). YME1L antibody used was a rabbit polyclonal antibody (1:1000, AP4882a) from Abgent (Suzhou city, Jiangsu Province, China). Opa1 antibody used was a mouse monoclonal antibody (1:1000, 18/Opa-1) from BD Biosciences (Franklin Lakes, NJ, United States). For loading controls, β-actin mouse monoclonal antibody (1:5000, 8H10D10) from Cell Signaling Technology (Beverly, MA, United States), β-tubulin rabbit polyclonal antibody (1:2500, T2200) from Millipore Sigma (St.Louis, MO, United States), and Cox IV rabbit polyclonal antibody (1:1000, 4844) from Cell Signaling Technology (Beverly, MA, United States) were used. The secondary antibodies used in this study include IRDye 680RD goat anti-rabbit and IRDye 800CW goat anti-mouse secondary antibodies, both at 1:15,000 dilution and obtained from Li-cor (Lincoln, NE, United States). Imaging and quantification was performed using the Li-Cor Odyssey Imaging System (Lincoln, NE, United States).

### Electron Microscopy

Fresh 1 cm^3^ pieces of tissue were fixed in 2.5% glutaraldehyde, 4% paraformaldehyde, 0.02% picric acid in 0.1 M sodium cacodylate buffer overnight at 4°C, washed, and then post fixed with osmium tetroxide reduced with potassium ferricyanide for 1 hour at room temperature, washed, and dehydrated in graded ethanol. Blocks generated using Embed 812 (Electron Microscopy Sciences, Hatfield, PA) resin were cut at 65 nm and mounted on copper grids, stained with uranyl acetate and lead citrate, and imaged using the JEM-1400 electron microscope (JEOL, Tokyo, Japan) as described previously. ^56^

### Statistics

Exploratory simple logistic regression models were fit to determine the optimal cut-off point for urine PHB and normalized PHB, which was the ratio of PHB and urine concentrations of creatinine, at which the odds of having relapse was maximized. We also used the Youden index, aimed at maximizing the sensitivity and specificity simultaneously, to find an optimal cut-off point. The Youden index is the maximum of (sensitivity + specificity − 1) over all threshold values of PHB and normalized PHB and allows evaluation of a diagnostic test with respect to its true positive and true negative rates. ^57^

Based on the results of these two criteria, two series of logistic regression models were used to evaluate whether PHB and normalized PHB (dichotomized by the optimal cut-off point) could predict Wilms tumor relapse. The first series was performed for the entire sample, considering either of the two PHB variables only. The second series were performed for patients with Wilms tumor. In addition to the PHB variables, three possible confounders were also included in the multivariate logistic regression models: Stage (I, II, or III), Histology (Blastema, Favorable Histology Wilms, Mixed Cell Wilms, or Epithelial), and Therapy (None vs. Any Chemotherapy). Backward selection was used to determine the most parsimonious model.

Fisher’s Exact tests were also applied to assess if the PHB or normalized PHB values categorized by the optimal cut-off point differed significantly among the three groups and the different relapse sites.

All statistical analyses and figures were performed using SAS version 9.2 (Cary, NC, United States), Origin version 9.1 (Northampton, MA, United States), and Graph Pad version 7.01 (LaJolla, CA, United States).

## Author Contributions

MO contributed to the study design, data collection, experiments, and interpretation of results. AK, SA, MB, AH, IM, S. Gunasekera, LG, GB, and JR performed experiments. TH, YH, AN, YYC, PC, and S. Gadd assisted with data analysis. MLQ, TH, JD, EP, EM, HS contributed to the interpretation of results. EM, HS, and AK contributed to the study design, data analysis, and interpretation. AK and MO wrote the manuscript, with contributions from other authors.

## Acknowledgements

We thank the Children’s Oncology Group Renal Tumor Committee for their support of this project. This study was supported by the Harvard Catalyst, Rally Foundation, Pablove Foundation, Hyundai Hope on Wheels, the Pediatric Cancer Research Foundation, Family and Friends of Caroline Bhatt, the Kristen Ann Carr Fund, Met Life Foundation, and the NCI K12 CA184746, R01 CA214812, U10 CA180899, and P30 CA008748. AK acknowledges the support of the St. Baldrick’s Foundation, Damon Runyon-Richard Lumsden Foundation, Josie Robertson Investigator Program, Burroughs Wellcome Fund, Cycle for Survival, and the Rita Allen Foundation. We thank Lee Cohen-Gould and Juan Pablo Jimenez for technical assistance, and Joseph Olechnowicz for comments on the manuscript.

## Notes

Conflict of interest: AK is a consultant for Novartis. The other authors have declared that no conflict of interest exists.

