## Supplemental Data for "Prohibitin is a prognostic marker of relapse and therapeutic target to block chemotherapy resistance in Wilms tumor"

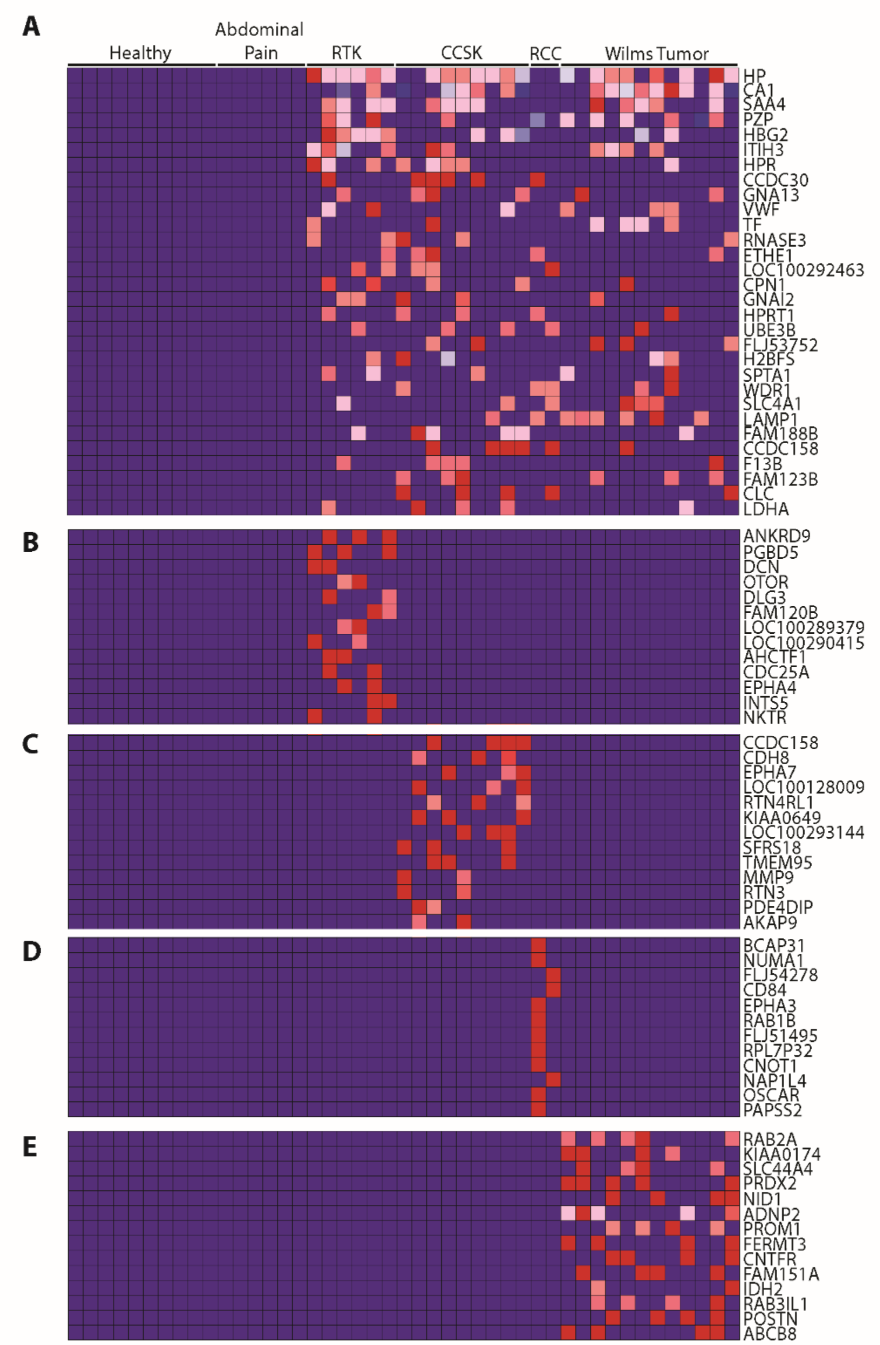


**Supplemental Figure 1. Using high accuracy mass spectrometry to profile the urine proteomes of childhood kidney tumors reveals markers of tissue injury and hematuria.**

(**A**). The 30 proteins which were most highly enriched in kidney tumors as compared with healthy controls and children with abdominal pain; RTK = Rhabdoid tumor of the kidney, CCSK = Clear cell sarcoma of the kidney, RCC = Renal cell carcinoma.

(**B-E**). The most highly enriched proteins in the urine of patients with rhabdoid tumors of the kidney (B), clear cell sarcomas of the kidney (C), renal cell carcinomas (D), and Wilms tumors (E) as compared with other pediatric renal tumors and controls.

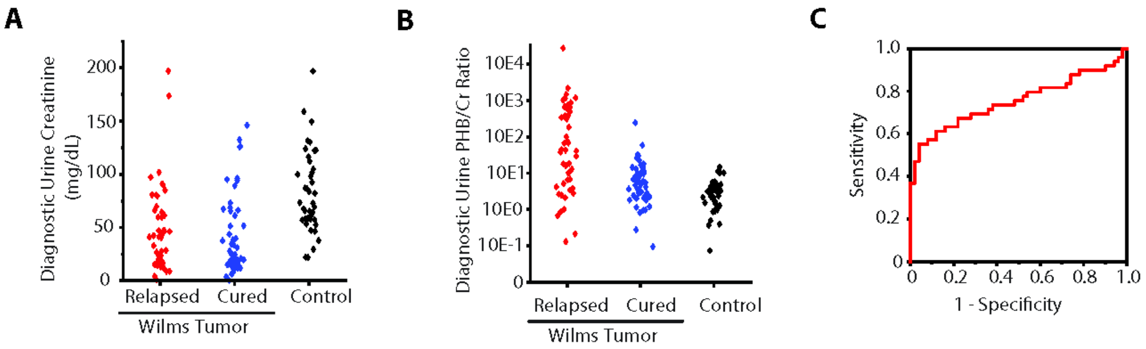


**Supplemental Figure 2. Elevated urine PHB/Cr ratio at diagnosis is a specific biomarker of relapse in favorable histology Wilms tumors.**

(**A-B**). Diagnostic urine creatinine levels (mg/dL) (A) and PHB/creatinine (Cr) levels (B) in favorable histology Wilms tumor patients who relapsed (Red, n = 49) are compared with those who were cured (Blue, n = 50) and normal controls (Black, n = 40).

(**C**). A receiver operating characteristic curve demonstrates the prognostic power of diagnostic urine prohibitin to predict relapse in favorable histology Wilms tumors at different sensitivity and specificity with an area under the curve of 0.75 (95% confidence interval 0.65-0.85).

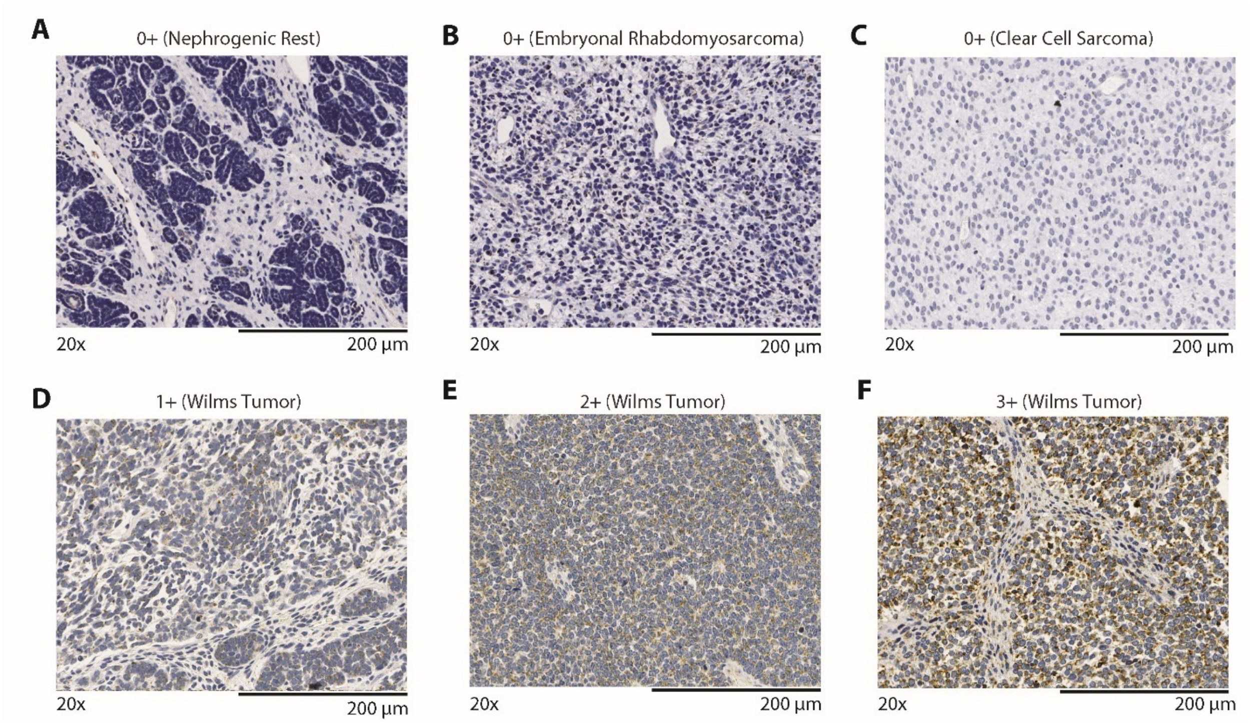


**Supplemental Figure 3. Tissue microarray immunohistochemistry standards and non-Wilms tumor controls**

The tissue microarray included several control tissues which did not express any PHB as shown. These were scored as 0+.

(**A**). Pre-malignant nephrogenic rests did not demonstrate PHB staining via IHC.

(**B**). Embryonal Rhabdomyosarcoma did not demonstrate PHB staining via IHC.

(**C**). Clear cell sarcoma did not demonstrate PHB staining via IHC.

(**D**). Wilms tumor tissues with low but evident expression of PHB were scored a 1+.

(**E**). Wilms tumor tissues with moderate expression of PHB were scored a 2+.

(**F**). Wilms tumor tissues with high expression of PHB were scored a 3+.

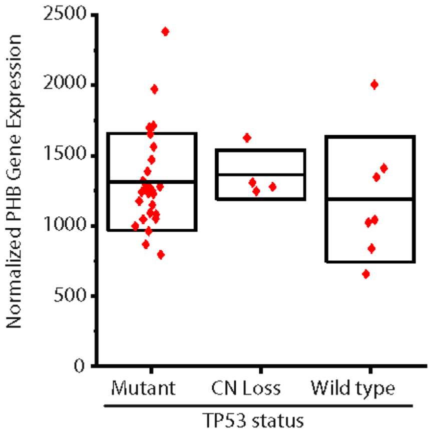


**Supplemental Figure 4. PHB mRNA expression does not correlate with *TP53* gene status in Wilms tumors with diffuse anaplasia**

Normalized PHB mRNA expression is compared to *TP53* gene status, separated by those with *TP53* mutations (N = 27), copy number (CN) loss (N = 4), or wild type (N = 7) in 38 diffusely anaplastic Wilms Tumors. Box blots overlaid in black demonstrate median and standard deviation.

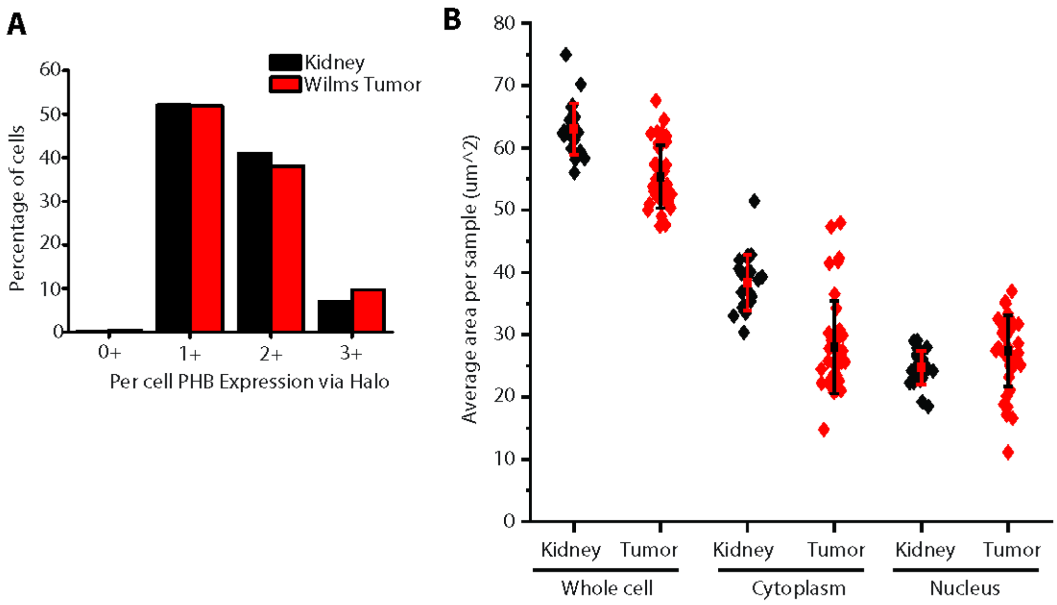


**Supplemental Figure 5. PHB expression in Wilms tumors compared with adjacent normal kidney tissues.**

(**A**). Percentage of cells with 0+ to 3+ PHB expression as measured by Halo imaging analysis software.

(**B**). A comparison of the average area of normal kidney (black diamonds) and Wilms tumor (red diamonds) whole cells, cytoplasm, and nuclei with overlaid median and standard deviations shown.

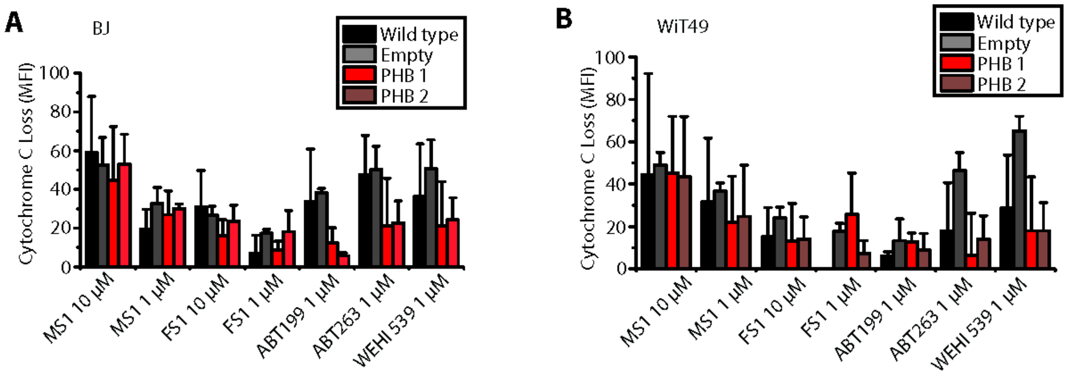


**Supplemental Figure 6. BH3 profiling reveals decreased apoptotic priming with PHB overexpression**

(**A-B**). Cytochrome c loss in response to treatment with different pro-apoptotic peptides comparing Wild type (Black), with Empty (Gray), and PHB overexpressing cells (Red, Brown) in BJ control fibroblast cells (A) and WiT49 Wilms tumor cells (B).

**Supplemental Table 1.** Characteristics of Wilms tumor patients in the validation ELISA cohort

| | | **Cured WT** | **Relapsed WT** |
|---|---|---|---|
| **Patients** | N | 50 | 49 |
| **Gender** (Male) | N (%) | 17 (34%) | 24 (49%) |
| **Age at diagnosis** (in Months) | Median (Range) | 43 (6-217) | 52 (5-129) |
| **Histology** | Favorable (%) | 50 (100%) | 49 (100%) |
| | Anaplastic (%) | 0 (0%) | 0 (0%) |
| **Stage** | I (%) | 8 (16%) | 4 (8%) |
| | II (%) | 18 (36%) | 22 (45%) |
| | III (%) | 24 (48%) | 23 (47%) |
| | IV (%) | 0 (0%) | 0 (0%) |
| | V (%) | 0 (0%) | 0 (0%) |
| **Chemotherapy** | Vincristine, Dactinomycin (%) | 23 (46%) | 23 (45%) |
| | Vincristine, Dactinomycin, Doxorubicin (%) | 25 (50%) | 23 (47%) |
| | Other/Unknown (%) | 2 (4%) | 4 (8%) |

**Supplemental Table 2**. Characteristics of Wilms tumor patients in the tissue microarray cohort

| **Patients** | N | 59 |
|---|---|---|
| **Gender** (Male) | N (%) | 26 (44%) |
| **Age at diagnosis** (in Months) | Median (Range) | 43 (6-120) |
| **Histology** | Favorable (%) | 59 (100%) |
| | Anaplastic (%) | 0 (0%) |
| **Stage** | I (%) | 11 (19%) |
| | II (%) | 19 (32%) |
| | III (%) | 20 (34%) |
| | IV (%) | 6 (10%) |
| | V (%) | 3 (5%) |
| **Chemotherapy** | Vincristine, Dactinomycin (%) | 0 (0%) |
| | Vincristine, Dactinomycin, Doxorubicin (%) | 0 (0%) |
| | Other/Unknown (%) | 59 (100%) |

**Supplemental Table 3.** Characteristics of Wilms tumor patients in the Halo analysis cohort

| **Patients** | N | 38 |
|---|---|---|
| **Gender** (Male) | N (%) | 14 (37%) |
| **Age at diagnosis** (in Months) | Median (Range) | 50 (10-236) |
| **Histology** | Favorable (%) | 23 (61%) |
| | Anaplastic (%) | 15 (39%) |
| **Stage*** | I (%) | 3 (11%) |
| | II (%) | 5 (18%) |
| | III (%) | 6 (21%) |
| | IV (%) | 11 (39%) |
| | V (%) | 3 (11%) |
| **Chemotherapy*** | Vincristine, Dactinomycin (%) | 8 (29%) |
| | Vincristine, Dactinomycin, Doxorubicin (%) | 16 (57%) |
| | Other/Unknown (%) | 4 (14%) |

*10 patients with limited clinical data available, so the stage and chemotherapy data only includes 28 patients for whom this data was available.
